# Analysis of cell-associated DENV RNA by oligo(dT) primed 5’ capture scRNAseq

**DOI:** 10.1101/2020.01.23.917757

**Authors:** Mark Sanborn, Tao Li, Kaitlin Victor, Hayden Siegfried, Christian Fung, Alan L. Rothman, Anon Srikiatkhachorn, Stefan Fernandez, Damon Ellison, Richard G. Jarman, Heather Friberg, Irina Maljkovic Berry, Jeffrey R. Currier, Adam T Waickman

## Abstract

Dengue is one of the most widespread vector-borne viral diseases in the world. However, the size, heterogeneity, and temporal dynamics of the cell-associated viral reservoir during acute dengue virus (DENV) infection remains unclear. In this study, we analyzed cells infected *in vitro* with DENV and PBMC from an individual experiencing a natural DENV infection utilizing 5’ capture single cell RNA sequencing (scRNAseq). Both positive- and negative-sense DENV RNA was detected in reactions containing either an oligo(dT) primer alone, or in reactions supplemented with a DENV-specific primer. The addition of a DENV-specific primer did not increase the total amount of DENV RNA captured or the fraction of cells identified as containing DENV RNA. However, inclusion of a DENV-specific cDNA primer did increase the viral genome coverage immediately 5’ to the primer binding site. Furthermore, while the majority of intracellular DENV sequence captured in this analysis mapped to the 5’ end of the viral genome, distinct patterns of enhanced coverage within the DENV polyprotein coding region were observed. The 5’ capture scRNAseq analysis of PBMC not only recapitulated previously published reports by detecting virally infected memory and naïve B cells, but also identified cell-associated genomic variants not observed in contemporaneous serum samples. These results demonstrate that oligo(dT) primed 5’ capture scRNAseq can detect DENV RNA and quantify virus-infected cells in physiologically relevant conditions, and provides insight into viral sequence variability within infected cells.

**IMPORTANCE:** Dengue is one of the most widespread vector-borne viral diseases in the world. However, it is still unclear which cells harbor virus during DENV infection, and how viral reservoirs in serum and infected cells are related. These results demonstrate for the first time that intracellular DENV RNA can be identified and infected cells quantified by 5’ capture scRNAseq. This strategy provides a significantly greater throughput and higher sensitivity than previously published methods and has the potential to provide additional information on the genomic heterogeneity of intracellular DENV RNA.

## INTRODUCTION

Dengue is one of the most widespread vector-borne viral diseases in the tropical and sub-tropical regions of the world. The causative agent– dengue virus (DENV) – is a positive-stranded RNA virus maintained in an anthroponotic cycle between the *Aedes aegypti* mosquito and humans [1]. Consisting of four co-circulating but genetically and immunologically distinct serotypes (DENV-1, -2, -3, and -4), DENV is thought to infect between 280 and 550 million people worldwide every year [2, 3]. Although the majority of DENV infections are subclinical, as many as 100 million infections every year result in symptomatic dengue fever. In addition, up to 500,000 infections per year result in severe dengue, which has a mortality rate of nearly 2.5% [4-7].

Following introduction into a human host by an infected mosquito during a blood meal acquisition, DENV asymptomatically replicates for 3-14 days prior to the onset of viremia or any clinical manifestation of infection [8]. After a presumed initial round of replication at the site of infection within tissue-resident or tissue-transiting leukocytes, DENV has been thought to disseminate and replicate within phagocytic mononucleocytes such as dendritic cells, monocytes, and macrophages which express the surface receptors DC-SIGN and/or mannose receptor [9-12]. However, recent studies utilizing techniques such as flow cytometry, RNAseq, and quantitative RT-PCR have demonstrated that B cells represent the major circulating cellular reservoir of DENV in individuals experiencing a natural DENV infection [13-15]. In any case, quantifying the cell-associated viral burden of DENV has the potential to provide actionable information in the setting of acute dengue, as differences in the cellular tropism/burden of DENV has been demonstrated in at least one report to correlate with the clinical severity of infection and with previous dengue exposure [13].

Recent advances in single cell RNA sequencing (scRNAseq) technology have revolutionized the field of cellular biology, providing insight into the heterogeneity of cellular transcription in an unbiased yet high-resolution fashion [16]. scRNAseq has also been leveraged to quantify the cellular tropism of several RNA viruses including influenza [17, 18], West Nile [19], Zika [20], and DENV [20, 21]. The majority of these published reports utilized a variant of the Smart-seq2 scRNAseq technology, wherein individual cells are deposited into separate wells in a 96 well plate containing the necessary reagents for cDNA synthesis and mRNA barcoding [22]. In addition to an oligo(dT) primer used to capture mammalian mRNA, these studies utilize a custom pathogen-specific primer during the cDNA synthesis reaction to maximize viral RNA recovery [20, 21]. While the published DENV-targeted Smart-seq2 methodology for DENV has demonstrated the potential to provide full-length viral sequence information, there are several limitations to the approach that may impede its broader adoption. Firstly, the targeted Smart-seq2 approach is low-throughput and relatively labor intensive even with modern fluid-handling robotics. Secondly, relying on a targeted primer for the detection and quantification of DENV RNA leaves open the possibility that divergent viral species will not be sufficiently primed to allow for downstream quantification.

An alternative to the commonly used Smart-seq2 scRNAseq methodology is 5’ capture scRNAseq, wherein only the 5’ end of a transcript is captured in the final sequencing library and tagged in such a manner to allow for cell-specific deconvolution [16, 23]. While this approach theoretically only captures the 5’ end of any transcript primed by the proffered cDNA synthesis primer (conventionally an oligo(dT) primer), it has the significant advantage of being compatible with several massively-parallel microfluidics-based platforms that allow for the simultaneous analysis of thousands of cells in a single reaction [16]. Although 5’ capture technology does not theoretically afford the same transcript coverage as other techniques, the total transcript recovery and unique gene recovery rate per cell is comparable with other technologies, and the exponentially greater number of cells analyzed allows for an unparalleled level of population resolution [16, 23]. Therefore, given the theoretical advantages offered by a 5’ capture approach, we attempted to determine if the technology is compatible with cell-associated DENV detection and quantification in the presence or absence of a virus-specific cDNA synthesis primer.

In this study, we first analyzed cells infected *in vitro* with DENV-1 utilizing 5’ capture scRNAseq. Despite the absence of a canonical polyadenylated tail, DENV RNA was incorporated in the 5’ capture scRNAseq reactions containing only an oligo(dT) primer, as well as in reactions supplemented with a DENV-specific primer. The addition of a DENV-specific cDNA synthesis primer did not appreciably increase either the total amount of DENV RNA captured by 5’capture scRNAseq analysis or the fraction of cells flagged as containing DENV RNA. The only effect observed upon the addition of a DENV-specific primer was an increase in the viral genome coverage proximal to the primer binding site. The majority of the DENV sequences recovered aligned to the 5’ end of the viral genome, but a regular pattern of higher-density coverage was noted across all *in vitro* infection samples. We then verified that 5’ capture scRNAseq can be utilized to detect/quantify cell-associated DENV RNA in PBMC from an individual experiencing a natural secondary DENV-1 infection. This analysis recapitulated previous reports of DENV-infected B cells and identified cell-associated viral variants that were not present in contemporaneous serum samples. These results demonstrate for the first time that DENV can be detected and quantified in physiologically relevant conditions using oligo(dT) primed 5’ capture scRNAseq, and provides insight into sequence variability of the DENV genome contained within infected cells.

## RESULTS

### Detection of DENV RNA by 5’ capture scRNAseq

To test the feasibility of quantifying intracellular DENV RNA in a 5’ capture scRNAseq assay - and the necessity for a DENV-specific primer in the RT reaction – we first utilized an *in vitro* cell infection model (**Figure 1A**). A DC-SIGN (CD209) expressing CEM.NK^R^ cell line (NKR2001A) was infected with DENV-1 (strain Nauru/West Pac/1974) at a MOI of 1 and cultured for 18 hours prior to harvest and analysis. Surface DENV-1 NS1 expression was assessed to confirm productive infection, and NS1^pos^ cells were sorted from the DENV-1 infected culture (**Figure 1B**). The uninfected parental CEM.NK^R^.DC-SIGN cell line, the bulk-DENV1 infected culture, and the sorted DENV-1 infected NS1^pos^ population were all subjected to scRNAseq 5’ capture analysis using a standard oligo(dT) RT primer for cDNA synthesis (**Supplemental Figure 1**). In addition, a parallel scRNAseq 5’ capture assay was performed on the sorted DENV1 infected NS1^pos^ population where a DENV-specific primer was added in addition to the conventional oligo(dT) primer at a 1:100 ratio (**Supplemental Figure 1**). Illumina-compatible NGS libraries were then generated from all samples, and sequenced to an equivalent depth on a NovaSeq6000 instrument with a 2×150bp paired-end (PE) read configuration. Samples were subsequently demultiplexed and aligned against the human GRCh38 or DENV-1(NCBI Reference Sequence NC_001477.1) reference genomes.

**Figure 1.**
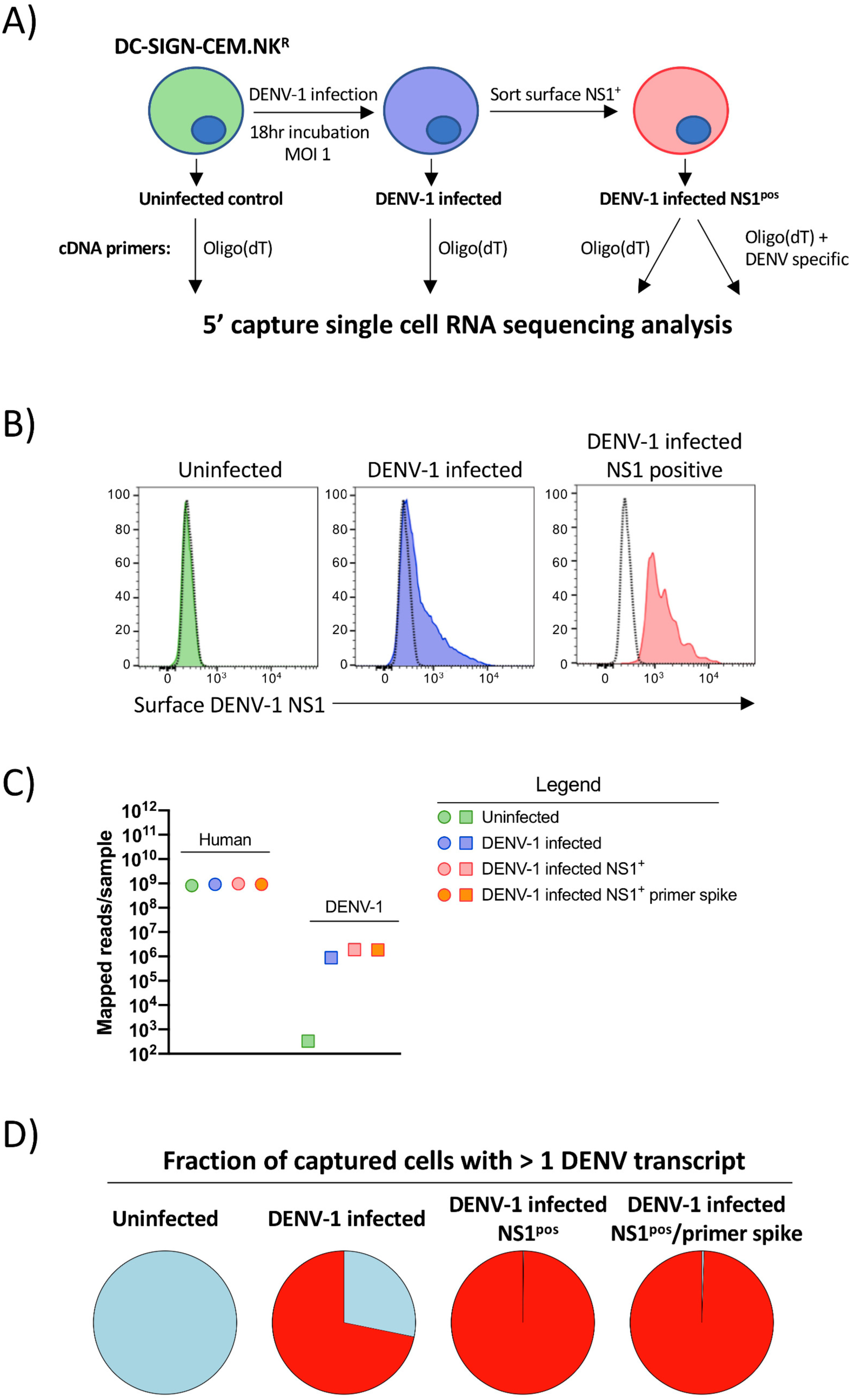
5’ capture scRNAseq analysis of *in vitro* DENV infected cells. **A)** Schematic representation of the infection, isolation, and analysis strategy utilized in this study. **B)** Surface DENV-1 NS1 expression on uninfected, DENV-1 infected, and sorted NS1^pos^ DENV-1 infected DC-SIGN expressing CEM.NK^R^ cells utilized for scRNAseq analysis. **C)** Number of PE reads confidently mapped from each sample to either the Gh38 human genome assembly or the DENV-1 (strain WestPac74) genome from each sample **D)** Fraction of cells from each sample with >=1 confidently mapped DENV-1 transcript present

An equivalent number of PE reads from all samples were confidently aligned and annotated against the human GRCh38 reference genome, indicating that neither DENV-1 infection nor the addition of the DENV-specific primer to the cDNA synthesis reaction significantly impacted the read depth/quality of the host genome-derived RNA sequence data (**Figure 1C, Table 1)**. In contrast, the number of PE reads confidently mapped against the DENV-1 reference genome varied significantly between the uninfected and DENV-infected samples, with the least number of reads identified in the uninfected samples (334 of 7.2×10^8^ total PE reads), and the most in the sorted NS1^pos^ DENV1 infected sample (1.9×10^6^ of 9.6×10^8^ total PE reads). The number of reads confidently mapped to the DENV-1 reference genome was additionally reflected in the fraction of captured cells flagged as containing >=1 transcript of DENV RNA (**Figure 1D**). No cells from the uninfected cell culture dataset were flagged as containing DENV RNA, while ∼70% of the cells from the bulk DENV infected culture were identified as harboring DENV RNA. Furthermore, more than 98% of the cells in both of the sorted NS1^pos^ DENV-1 infected samples were identified as containing DENV-1 RNA. These data demonstrate that DENV RNA can be detected using 5’ capture scRNAseq analysis, and that a DENV-specific primer is not necessary to obtain this result.

**Table 1.** Sequencing and annotation statistics for *in vitro* infection experiments.

| Sample name | Total reads in sample | Reads confidently mapped to Genome | Median unique genes detected per cell | Median UMI counts per cell | Total genes detected in sample |
| --- | --- | --- | --- | --- | --- |
| Uninfected | 364,378,131 | 84.7% | 5,680 | 40,998 | 17,796 |
| DENV-1 infected | 436,057,561 | 84.7% | 5,416 | 37,282 | 18,410 |
| DENV-1 infected NS1<sup>pos</sup> | 478,167,079 | 86.1% | 5,530 | 40,664 | 18,719 |
| DENV-1 infected NS1<sup>pos</sup> + DENV1 primer | 443,086,874 | 88.7% | 5,507 | 39,951 | 18,610 |

### Quantification of DENV RNA by 5’ capture scRNAseq

To better characterize the DENV RNA identified in our scRNAseq 5’ capture assay, we examined the abundance and length of each unique RNA template tagged with a unique molecular identifier (UMI) that mapped to the DENV1 reference genome. The number of unique UMI-tagged DENV RNA molecules identified (DENV RNA-positive) per cell ranged between 1 and 500 in all infected samples, with the sorted NS1^pos^ populations containing on average 2X the number of DENV-1 mapped UMIs as the unsorted DENV1 infected sample (**Figure 2A**). No significant increase in the number of UMIs/cell was observed within the sample containing the additional DENV specific primer. The length of the DENV-1 reads ranged between 100 and 600bp, which is consistent with the library preparation strategy utilized during the assay (**Figure 2B**). Correspondingly, on average each DENV-mapped UMI reflected ∼3% of the DENV-1 genome (**Figure 2C**). When aligned in aggregate, the total DENV genome recovered per cell ranged between 3% and 99%, with the median genome recovery rate significantly higher in the sorted NS1^pos^ DENV infected samples than in the bulk-infected sample (**Figure 2D**). On average, 50% full-length DENV-1 genome coverage was achieved with 38.3 UMIs per cell across all infected samples, while 75% genome coverage was achieved with 100 UMIs per cell (**Figure 2E**). Therefore, despite a single UMI providing a relatively small fragment of the DENV genome, a full-length cell-associated consensus genome can be constructed from relatively few unique transcripts.

**Figure 2.**
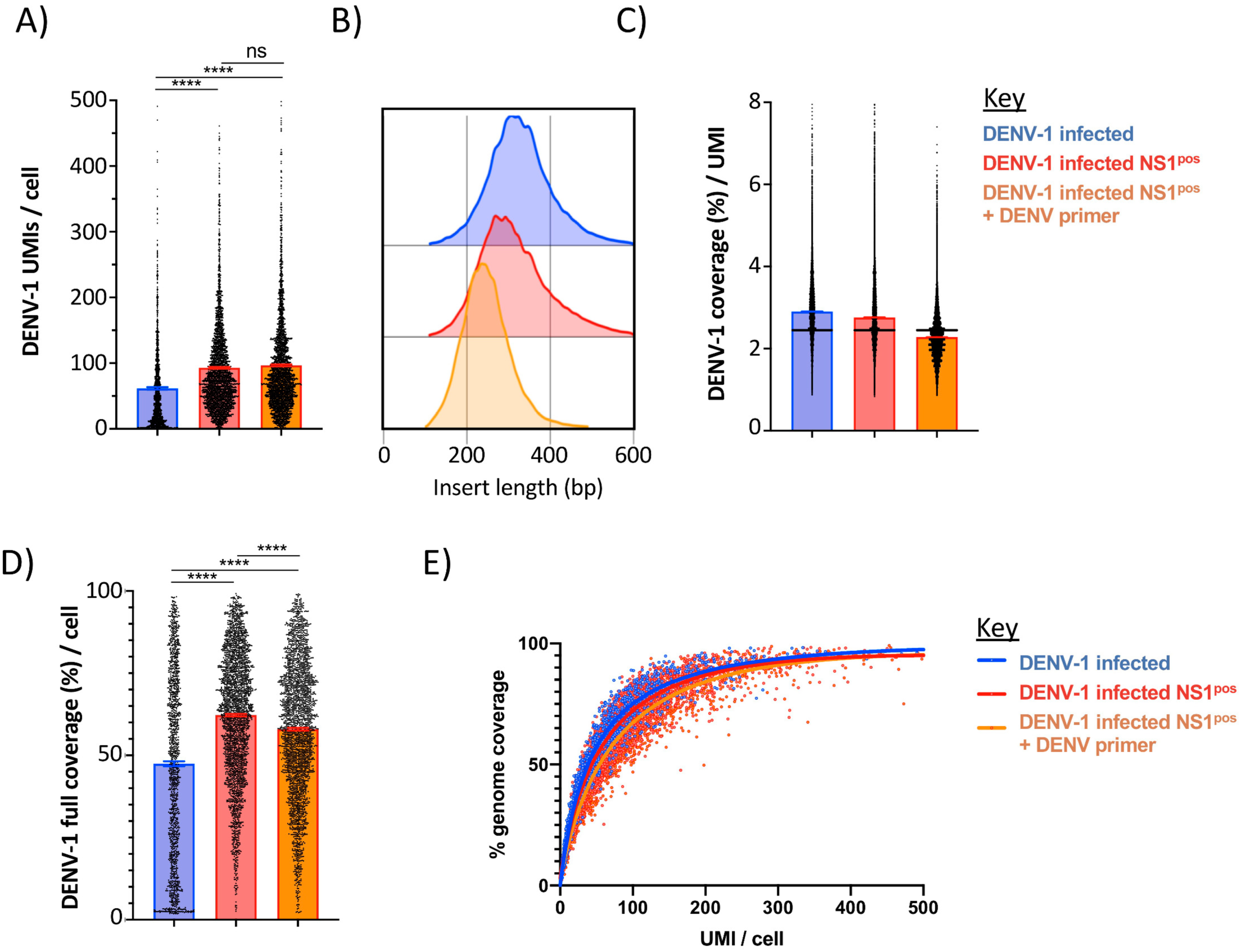
Characteristics of DENV-1 mapped transcripts identified by 5’ capture scRNAseq. **A)** Number of unique DENV-1 mapped transcripts identified per cell in DENV-1 infected DC-SIGN expressing CEM.NK^R^ cell cultures (blue), DENV-1 infected sorted NS1^pos^ cultures (red), and DENV-1 infected sorted NS1 ^pos^ cultures supplemented with a DENV-specific cDNA synthesis primer during scRNAseq analysis (orange). **B)** DENV-1 mapped insert lengths identified by scRNAseq analysis **C)** Median DENV-1 genome coverage obtained per UMI in 5’ capture scRNAseq analysis **D)** DENV-1 genome coverage achieved per cell by 5’ capture scRNAseq. Only cells with >=1 DENV-1 mapped UMI are shown. **E)** Relationship between DENV-1 UMI recovery and DENV-1 genome coverage achieved by 5’ capture scRNAseq. Curves calculated by non-linear regression (single variable). **** p <0.0001, unpaired t test.

### Genomic coverage of positive-sense DENV generated by 5’ capture scRNAseq

Many cells captured in this analysis contained DENV RNA that confidently mapped across a significant fraction of the DENV-1 genome. Therefore, we attempted to ascertain whether there were regions of preferential sequence recovery in our analysis or if the recovery was uniform across the DENV genome. We additionally attempted to determine if the inclusion of a DENV-specific RT primer altered the distribution of DENV genomic sequence recovery.

Consistent with the 5’ capture technology utilized in this analysis, ∼27.5% to 32.0 of all positive-sense DENV-1 reads aligned to the first 500bp of the DENV genome in each sample (**Figure 3A, 3B, 3C**). However, a consistent pattern of enriched positive-sense genomic sequence recovery was noted in all samples within the 3’ end of the polyprotein encoding region of the DENV genome (**Figure 3A, 3B, 3C**). Inclusion of the DENV-specific primer increased the genomic coverage 300-500bp upstream of the primer binding site in (**Figure 3C**).

**Figure 3.**
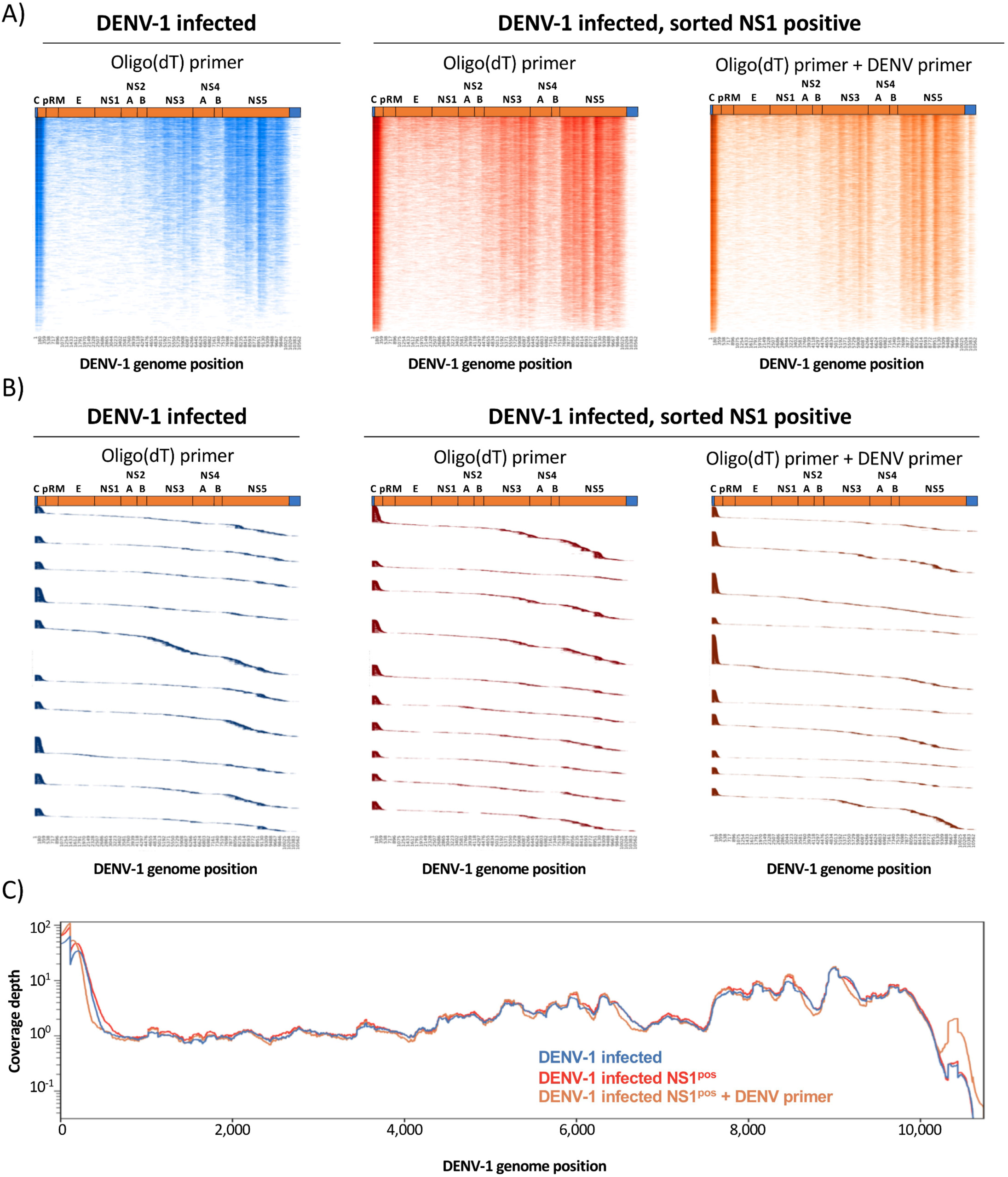
Genome coverage of positive-sense DENV-1 mapped transcripts identified by 5’ capture scRNAseq. **A)** Heatmap representation of positive-sense DENV-1 genomic sequence recovery in individual cells analyzed by 5’ capture scRNAseq from DENV-1 infected cultures (blue), DENV-1 infected sorted NS1^pos^ cultures (red), and DENV-1 infected sorted NS1^pos^ cultures supplemented with a DENV-specific cDNA synthesis primer (orange). Color intensity indicates sequence coverage/depth at that genomic location, vertical axis contains a single cell per line. Cells are ranked by total DENV-1 genomic coverage. Cells with >=1 DENV-1 mapped UMI are included in the visualization. **B)** Positive-sense DENV-1 genomic localization of individual UMIs obtained from the top 10 cells in each analysis condition with the most DENV-1 coverage as assessed by 5’ capture scRNAseq analysis. UMIs are ranked by genomic location 5’ to 3’, and the vertical axis contains a single DENV-1 mapped UMI per line. **C)** Aggregated positive-sense DENV-1 genomic coverage and sequence depth of all *in vitro* infected samples as assessed by 5’ capture scRNAseq

### Detection of negative-sense DENV RNA by 5’ capture scRNAseq

Productive DENV infection is accompanied by the generation of intracellular negative-sense DENV RNA [24]. As with other members of the virus family *Flaviviridae*, this intermediate antigenome is utilized as a template for the synthesis of additional positive-sense DENV genomic RNA [25]. In our analysis we observed that 67% of the unenriched DENV-1 infected CM.NK^R^ cells analyzed by oligo(dT) primed scRNAseq simultaneously contained reads that mapped to either the positive- or negative-sense orientation of the DENV genome (herein referred to as positive- or negative-sense DENV RNA, respectively) (**Figure 4A**). In contrast, 99% of sorted NS1^pos^ cells – which by definition are productively infected – contained both positive- and negative-sense DENV RNA (**Figure 4A**). Within DENV-infected cells, negative-sense DENV RNA accounted for ∼7.1% of all annotated DENV RNA in both unfractionated and sorted NS1^pos^ DENV-1 infected cells (**Figure 4B**). A linear relationship was observed between the amount of positive-sense DENV RNA identified within a cell and the corresponding abundance of negative-sense RNA (**Figure 4C**). However, the relatively low r^2^ value of these relationships (DENV-1 infected: 0.37. Sorted NS1^pos^ DENV-1 infected: 0.25) suggests that factors other than the abundance of positive-sense DENV contributes to the relative abundance of negative-sense DENV RNA within DENV infected cells.

**Figure 4.**
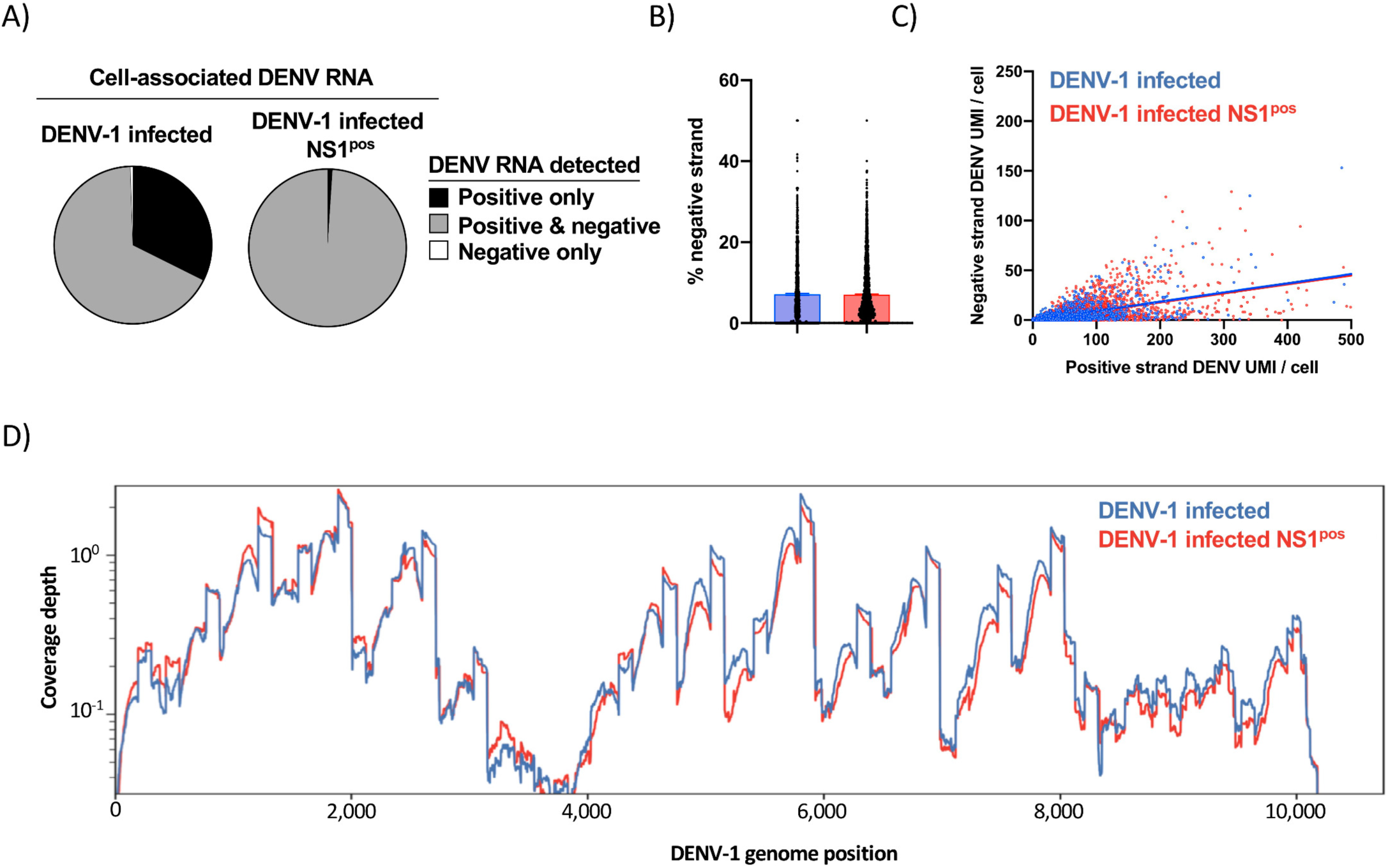
Characterization of negative-sense DENV RNA by 5’ capture scRNAseq. **A)** Fraction of indicated cell populations containing either only positive-sense DENV RNA (black), both positive- and negative-sense DENV RNA (gray), or only negative-sense DENV RNA (white) as quantified by 5’ capture scRNAseq. **B)** Percentage of all DENV-mapped RNA with a negative-sense orientation in DENV-1 infected (blue) or sorted NS1^pos^ DENV-1 infected cells as quantified by 5’ capture scRNAseq. **C)** Relationship between positive-sense and negative-sense DENV RNA abundance within DENV-1 infected cells. **D)** Aggregated negative-sense DENV-1 genomic coverage and sequence depth of *in vitro* infected samples as assessed by 5’ capture scRNAseq.

Enriched regions of negative-sense DENV RNA recovery was observed across the DENV genome in samples analyzed by 5’ capture scRNAseq (**Figure 4D**). However, the pattern of negative-sense DENV RNA recovery is distinct from that observed for positive-sense DENV RNA in the same samples. These results demonstrate that 5’ capture scRNAseq can identify negative-sense DENV RNA within DENV infected cells, and that the frequency of cells harboring negative-sense DENV RNA corresponds with the presumed frequency of productive DENV infection.

### Detection of DENV RNA in PBMC following natural infection by 5’ capture scRNAseq

To determine the feasibility of identifying and quantifying PBMC harboring DENV RNA following natural viral infection using 5’ capture scRNAseq, we analyzed PBMC samples obtained from an individual experiencing a natural secondary DENV1 infection. To maximize the accuracy and sensitivity of the 5’ capture scRNAseq dependent detection of DENV genomic RNA, a custom DENV-1 reference was created using the sequence of the serum-associated virus infecting the individual in this study (**Supplemental Figure 3, Table 2**). NGS analysis of the serum-associated DENV reservoir captured in this analysis provided a full-length genome with a coverage depth of 1000x or greater from both time points assessed in this study (fever day -2, fever day -1) (**Table 2**). A 20% variant level analysis showed no genomic sequence variants at either time point within the serum-associated viral reservoir. However, a 1% variant level analysis showed 4 called synonymous variants at position 445 and 9565 in the DENV polyprotein coding region that were present in equal frequencies at both time points (**Table 2**).

**Table 2:** Cell and serum associated DENV sequence variation.

| Genome Position | Fever Day -2 |  |  |  | Fever Day -1 |  |  |  |
| --- | --- | --- | --- | --- | --- | --- | --- | --- |
|  | Serum-associated |  | Cell-associated |  | Serum-associated |  | Cell-associated |  |
|  | Sequence Call, nt [AA] | Read Depth | Sequence Call, nt [AA] | Read Depth | Sequence Call, nt [AA] | Read Depth | Sequence Call, nt [AA] | Read Depth |
| 445 | 13.7% A [L]<br>86.3% G [L] | > 1,000 | 12.3% A [L]<br>87.7% G [L] | 187 | 13.1% A [L]<br>86.9% G [L] | > 1,000 | 10.5% A [L]<br>89.5% G [L] | 19 |
| 860 | C [H] | > 1,000 | 12.0% T [Y]<br>88.0% C [H] | 92 | C [H] | > 1,000 | C [H] | 55 |
| 1008 | T [L] | > 1,000 | 64.2% T [L]<br>35.8% C [P] | 81 | T [L] | > 1,000 | T [L] | 14 |
| 2576 | G [G] | > 1,000 | 32.0% T [C]<br>68.0% G [G] | 50 | G [G] | > 1,000 | G [G] | 17 |
| 5848 | T [I] | > 1,000 | T [I] | 174 | T [I] | > 1,000 | 78.9% T [I]<br>21.1% C [I] | 57 |
| 7042 | G [Q] | > 1,000 | G [Q] | 73 | G [Q] | > 1,000 | 32.7% A [Q]<br>67.3% G [Q] | 52 |
| 7662 | G [R] | > 1,000 | 14.1% A [K]<br>85.9% G [R] | 156 | G [R] | > 1,000 | G [R] | 24 |
| 9127 | C [L] | > 1,000 | 22.8% T [L]<br>77.2% C [L] | 79 | C [L] | > 1,000 | C [L] | 24 |
| 9276 | T [F] | > 1,000 | T [F] | 79 | T [F] | > 1,000 | 20.0% T [F]<br>80.0% C [S] | 15 |
| 9565 | 98.5% C [C]<br>1.5% T [C] | > 1,000 | C [C] | 25 | 98.4% C [C]<br>1.6% T [C] | > 1,000 | C [C] | 19 |

nt – nucleotide, AA- amino acid

Two sequential PBMC samples were analyzed from this individual during the acute stage of infection (fever day -2, fever day -1), as well as a ‘baseline’ sample from 180 days post defervescence. Total viable CD3^+^ T cells and CD19^+^ B cells were sorted from the cryopreserved sample and analyzed by 5’ capture scRNAseq using an oligo(dT) primer for cDNA synthesis. A total of 9,565 cells were captured across all 3 time points, and 11 transcriptionally distinct cell populations annotated using cell-associated transcripts confidently aligned against the GRCh38 reference genome (**Figure 5A, Supplemental Figure 4, Supplemental Table 1**). Cell-associated transcripts not mapped to the GrCh38 reference genome were subsequently aligned against a custom DENV-1 reference generated from the sequence of the serum-associated virus identified by targeted NGS sequencing (**Supplemental Figure 3, Table 2**). Consistent with previously published reports, DENV-1 transcripts were identified within the peripheral B cell compartment in both naïve and memory-phenotype B cells (**Figure 5B, 5C**). Both positive- and negative-sense DENV RNA was identified within the cells, indicating that they were productively infected (**Supplemental Figure 5**). Although the total amount of DENV RNA recovered in this assay was insufficient to provide a full-length genome in any individual cell (**Figure 5D, 5E**), in aggregate there was enough cell-associated DENV genomic material to generate a consensus genome sequence across both acute infection time points (**Figure 5F, Table 2**).

**Figure 5.**
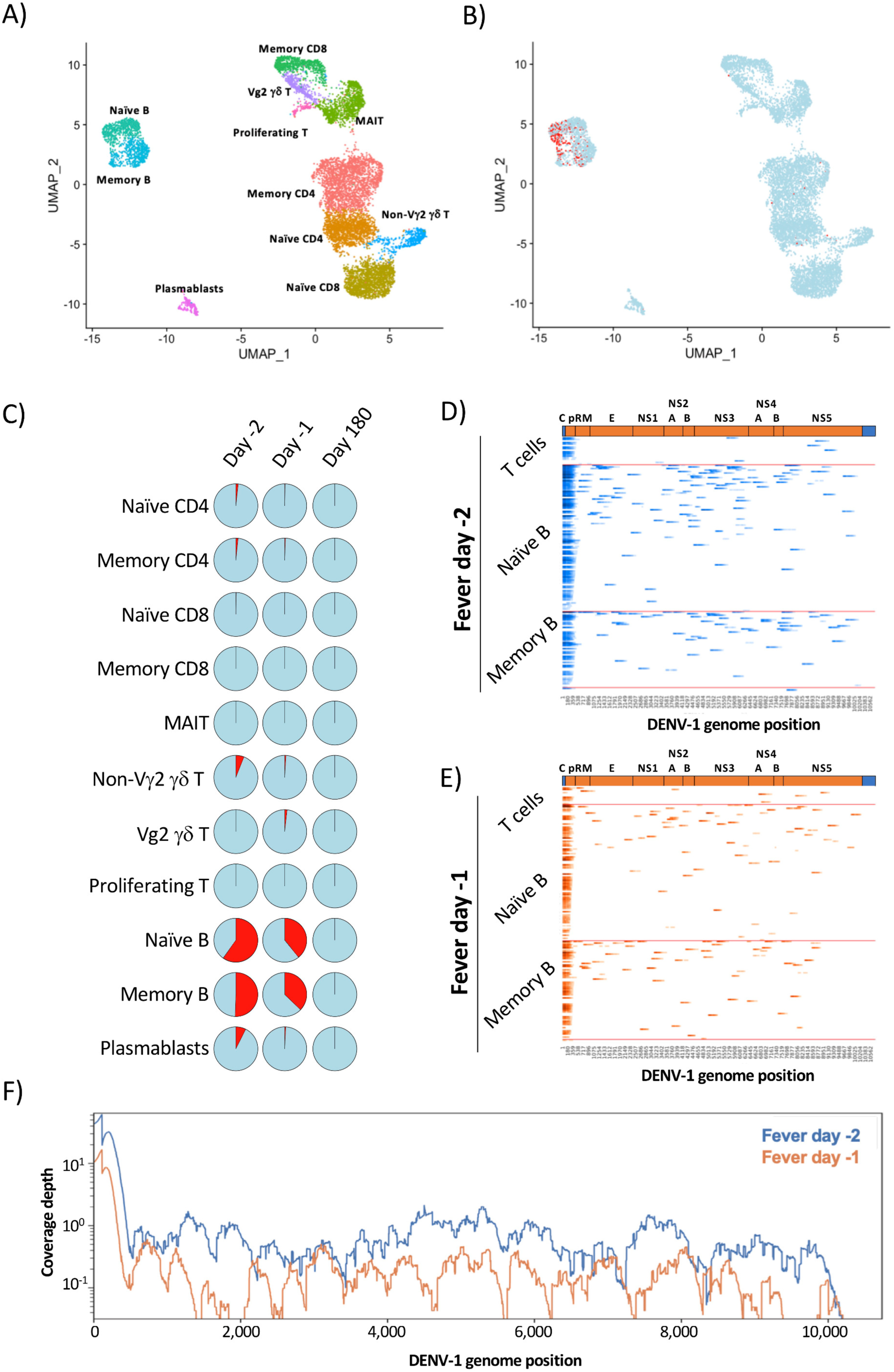
Identification and quantification of virally infected PBMC following natural DENV infection. **A)** UMAP projection and population level-annotation of 9,565 sorted CD3^+^ and CD19^+^ PBMC analyzed by 5’ capture scRNAseq. Plot is an aggregated analysis of 3 timepoints from the same DENV-1 infected individual (fever days -2, -1, +180) **B)** UMAP projection of sorted PBMCs captured by 5’capture scRNAseq indicating uninfected cells (blue) and cells containing >=1 DENV-1 mapped UMI (red) **C)** Proportion of the identified PBMC populations from each analyzed time point containing >=1 UMI confidently mapped to the DENV-1 genome **D)** Heatmap representation of DENV-1 genomic sequence recovery in individual cells analyzed by 5’ capture scRNAseq from the fever day -2 PBMC sample. Infected cells are binned by population identity. **E)** Heatmap representation of DENV-1 genomic sequence recovery in individual cells analyzed by 5’ capture scRNAseq from the fever day -1 PBMC sample. Infected cells are binned by population identity. **F)** Aggregated DENV-1 genomic coverage and sequence depth of all *in vivo* infected samples as assessed by 5’ capture scRNAseq

Consistent with the coverage results obtained from the *in vitro* infection experiments, the majority of the confidently mapped DENV-1 reads came from the first 500bp of the DENV-1 genome (Day -2: 50.73%. Day -1: 53.3%). Although the cell-associated DENV genome sequencing depth was limited in this analysis, the synonymous genomic variants observed at position 445 within the serum-associated viral reservoir were also observed within the cell-associated viral reservoir at a nearly identical frequency at both time points analyzed (**Table 2**). In addition, a further 16 variants at 8 genomic positions were observed within the cell-associated viral reservoir that were not observed within the contemporaneous serum-associated viral reservoirs (**Table 2**). While the intracellular DENV sequence depth is limited in this analysis, these results suggest a greater degree of genomic variation within the cell-associated DENV viral reservoir during acute DENV infection than in the serum-associated viral reservoir.

## DISCUSSION

In this study we demonstrate that DENV genomic RNA can be captured and quantified by 5’ capture scRNAseq. Despite the absence of a conventional polyadenylated 3’ tail, DENV genomic RNA was successfully captured utilizing an oligo(dT) primer without addition of a DENV-specific RT primer. The addition of a DENV-specific primer during the cDNA synthesis step did not appreciably increase the number of DENV RNA molecules captured by 5’ scRNAseq analysis. The majority of the DENV sequences recovered in this analysis aligned to the 5’ end of the viral genome, but a regular pattern of higher-density coverage was noted across all *in vitro* infection samples as well. As sequence recovery corresponds to locations where cDNA synthesis terminates in 5’ capture scRNAseq, this pattern may reflect unappreciated secondary structures in the native intracellular DENV genome that impeded the reverse transcriptase processivity. We additionally verified that 5’ capture scRNAseq can be utilized to detect/quantify physiologically-relevant levels of cell-associated DENV RNA by analyzing samples from an individual experiencing a natural secondary DENV-1 infection. This analysis recapitulated previous reports by detecting virally infected B cells, and, in addition, we were able to compare the sequence of the cell-associated and serum virus in the same individual. While both cell-associated and serum viruses possessed an identical intra-host single-nucleotide variant (iSNV) with approximately equal frequencies in one of the genomic positions, additional iSNVs were observed within the cell-associated viral reservoir that were not observed within the contemporaneous serum. While it is well known that DENV exists as a population of viral variants within a host, here for the first time we show that these viral populations are similar, though not identical, in composition and frequency within these two different host compartments. Virus population compartmentalization has been well described in HIV [26], and has been associated with disease severity. Whereas the chronic nature of HIV infection is in stark contrast to DENV, more studies are needed to determine if population compartmentalization and heterogeneity has any relationship to dengue severity.

The observation that DENV genomic RNA is captured in a reverse transcription reaction utilizing anchored-oligo(dT) priming was initially unexpected but is not without precedent. Previous studies have demonstrated that cDNA can be generated from flavivirus genomic RNA templates using an oligo(dT) primer, although the exact oligo(dT) primer binding sites were not resolved [27]. Furthermore, the anchored-oligo(dT) primer used in this analysis contains a degenerate 3’ terminus, which may increase the likelihood of the oligo binding/priming off a sequence other than a canonical mRNA poly-A tail. Indeed, internal transcript priming at A rich regions is a long-appreciated problem with oligo(dT) based RT-methods [28-32]. However, while it is plausible – even probable - that the oligo(dT) primer used in the analysis presented here bound to polypyrimidine tracts within the DENV genome, the exact location of these internal priming events cannot be determined using this approach. This is due to the fact that the 5’ capture assay used in this study preferentially captures sequences from regions of RT-termination, rather than RT-initiation [16, 23].

While the location of RT initiation cannot be resolved by 5’ scRNAseq, this approach may unexpectedly allow for the resolution of native secondary/tertiary structures within the intracellular DENV genome. Extensive secondary structures within an RNA template can severely impede the processivity of a reverse transcriptase, leading to the production of truncated cDNA molecules downstream of the initial priming event [33, 34]. This inherent feature of RT-dependent cDNA synthesis may be exacerbated by the fact that the MMLV-derived RT used in this 5’ capture assay displays maximal enzymatic activity at 43°C: a temperature at which significant secondary structures will be retained in single-stranded RNA templates. The observation that the inclusion of a DENV-specific RT primer only increases the depth of the recovered DENV genome at 200-500bp downstream of the primer binding site reinforces the concept that RT processivity overall is profoundly limited on the DENV genome in the described scRNAseq assay. Therefore, if RT processivity is indeed limited on the DENV genome in this assay, oligo(dT) priming must occur frequently in order to achieve the levels of full genome coverage observed in our analysis.

While secondary structures within the 5’ and 3’ UTRs of the DENV genome are critical for viral replication, propagation, and genomic stability [35-38], up to one third of the nucleotides in the DENV genome have been suggested to be involved in the formation of higher-order structures [39]. Conserved regions of secondary structure within the 5’ protein coding regions of the DENV-2 genome are known to play a role in regulating the translation of the DENV polyprotein [40], while hairpin structures in the capsid region of multiple flaviviruses are critical for the initiation of viral RNA synthesis [41, 42]. While significant secondary/tertiary structural features have been predicted to exist within the 3’ region of the DENV polyprotein-coding RNA sequence [39], the prevalence and biological role of these higher-order structures *in vivo* are currently unknown.

The ability to leverage massively parallel scRNAseq methods for the assessment of DENV cellular tropism represents an important technical advancement. By facilitating higher-resolution analysis of the cellular tropism, infection kinetics, and transcriptional consequences associated with DENV infection, this methodology may enable the identification of more accurate correlates of risk during acute DENV infection and potentially improve diagnostic accuracy.

## MATERIALS AND METHODS

### Cell lines

The parental CEM.NK^R^ cell line was obtained through the NIH AIDS Reagent Program, Division of AIDS, NIAID, NIH, courtesy of Dr. Peter Cresswell [43]. A stably transfected and cloned, DC-SIGN (CD209: Genbank NM_021155) expressing variant of this cell line (NKR2001A) was developed for DENV-infection experiments.

### Viruses

DENV1 (strain Nauru/West Pac/1974) was propagated in C6/36 mosquito cells was utilized for i*n vitro* infection assays. Virus was purified by ultracentrifugation through a 30% sucrose solution and the virus pellet was resuspended in PBS.

### Sample collection

Peripheral blood mononuclear cells (PBMC) were isolated from whole blood samples collected from children enrolled in a hospital-based acute illness study in Bangkok, Thailand [44, 45]. The study included children who presented to the hospital with suspected dengue. Blood draws were obtained serially during acute infection and at early and late convalescent time points; the term ‘fever day’ is used to report acute illness time points relative to Day 0, defined as the day of defervescence. The infecting virus type (DENV-1-4) was determined by virus isolation and/or RT-PCR [46], and serology (EIA and HAI assays) was used to distinguish primary and secondary DENV infections [47]. Written informed consent was obtained from each subject and/or his/her parent or guardian. The study protocol was approved by the Institutional Review Boards of the Thai Ministry of Public Health, the Office of the U.S. Army Surgeon General, and the University of Massachusetts Medical School. PBMC and plasma samples were cryopreserved for later analysis.

### Flow Cytometry and cell sorting

Cryopreserved PBMC were thawed and placed in RPMI 1640 medium supplemented with 10% heat-inactivated normal human serum (100–318, Gemini Bio-Products), L-glutamine, penicillin, and streptomycin prior to analysis. Cell viability was assessed by trypan blue exclusion. Surface staining for flow cytometry analysis and cell sorting was performed in PBS supplemented with 2% FBS at room temperature. Aqua Live/Dead (ThermoFisher, L34957) was used to exclude dead cells in all experiments. Antibodies and dilutions used for flow cytometry analysis are listed in **Supplemental Table 2**. Cell sorting was performed on a BD FACSAria Fusion instrument, and data analyzed using FlowJo v10.2 software (Treestar).

### Single-cell RNA sequencing library generation

Flow-sorted viable B and T cell suspensions or DC-SIGN expressing CEM.NK^R^ cells were prepared for single-cell RNA sequencing using the Chromium Single-Cell 5′ Reagent version 2 kit and Chromium Single-Cell Controller (10x Genomics, CA) [48]. Approximately 2000–8000 cells per reaction suspended at a density of 50–500 cells/μL in PBS plus 0.5% FBS were loaded for gel bead-in-emulsion (GEM) generation and barcoding. Reverse transcription, RT-cleanup, and cDNA amplification were performed to isolate and amplify cDNA for downstream 5′ gene or enriched V(D)J library construction according to the manufacturer’s protocol. Libraries were constructed using the Chromium Single-Cell 5′ reagent kit, 5′ Library Construction Kit, and i7 Multiplex Kit (10x Genomics, CA) according to the manufacturer’s protocol.

### Sequencing

scRNAseq 5′ gene expression libraries were sequenced on an Illumina NovaSeq 6000 instrument using the S1, S2, or S4 reagent kits (300 cycles). Libraries were balanced to allow for ∼150,000 reads/cell for 5′ gene expression libraries. Sequencing parameters were set for 150 cycles for Read1, 8 cycles for Index1, and 150 cycles for Read2. Prior to sequencing, library quality and concentration were assessed using an Agilent 4200 TapeStation with High Sensitivity D5000 ScreenTape Assay and Qubit Fluorometer (Thermo Fisher Scientific) with dsDNA BR assay kit according to the manufacturer’s recommendations

### 5’ host gene expression analysis/visualization

5′ gene expression alignment from *in vitro* infected DC-SIGN expressing CEM.NK^R^ cells or sorted PBMC was performed using the 10x Genomics Cell Ranger pipeline [48]. Sample demultiplexing, alignment, barcode/UMI filtering, and duplicate compression was performed using the Cell Ranger software package (10x Genomics, CA, v2.1.0) and bcl2fastq2 (Illumina, CA, v2.20) according to the manufacturer’s recommendations, using the default settings and mkfastq/count commands, respectively. Transcript alignment was performed against a human reference library generated using the Cell Ranger mkref command and the Ensembl GRCh38 v87 top-level genome FASTA and the corresponding Ensembl v87 gene GTF. Multi-sample integration, data normalization, visualization, and differential gene expression was performed using the R package Seurat (v3.0) [49, 50].

### 5’ DENV-1 genome analysis

Quantification and alignment of cell-associated DENV-1 genomic RNA from was performed using the 10x Genomics Cell Ranger pipeline [48]. Sample demultiplexing, alignment, barcode/UMI filtering, and duplicate compression was performed using the Cell Ranger software package (10x Genomics, CA, v2.1.0) and bcl2fastq2 (Illumina, CA, v2.20) according to the manufacturer’s recommendations, using the default settings and mkfastq/count commands, respectively. Transcript alignment was performed against a custom DENV-1 reference library generated using the Cell Ranger mkref command and the DENV-1 (strain Westpac74) FASTA and the corresponding gene GTF, or a custom DENV-1 FASTA generated from targeted NGS analysis of contemporaneous serum samples from the individual analyzed in the study.

### Cell associated DENV UMI dependent genomic visualization

DENV-1 mapped reads were provided in BAM format from the Cell Ranger pipeline. UMI-tools dedup was used to deduplicate the BAM files on a per cell basis using cell barcode, UMI barcode, and alignment position of the paired end reads [51]. BAM files were split into positive-sense and negative-sense mapped reads using BAM alignment tags provided by the Cellranger pipeline. Only read pairs where both reads mapped to the same genomic orientation were utilized in downstream visualizations. Using a list of cell barcodes identified as being associated with an intact cell by the cellranger count analysis of GRCh38 aligned transcripts, cell and unique UMI specific statistics were extracted using an in-house pipeline. In short, the BAM files were subset by cell barcode using 10x Genomics subset-bam followed conversion to fastq using bedtools bamToFastq [52]. The reads were then remapped using BWA-MEM and the BWA produced BAM files were analyzed using a custom Python script with a samtools [53] depth frontend. Unique UMI codes were extracted from the subset cell BAM and used to further subset the individual cell BAMs. The unique UMI assemblies were then analyzed similarly for assembly and coverage statistics. DENV-1 (strain Westpac74) was used as a reference for the *in vitro* samples and the constructed serum consensus was used for the natural infection samples.

In addition to statistical files the pipeline outputs pickled python dictionaries with cell barcode or UMI keys and list values (with length of the reference genome) of genome depth of coverage. These dictionaries were used in the heatmap visualization of DENV-1 coverage. Individual cell coverage was ordered by percent coverage and the maximum heatmap intensity was set at a threshold three standard deviations above the mean. Unique UMIs were plotted for the top 10 cells in each in-vitro sample. UMI were binned by cell and ordered by position in the DENV-1 genome. A low number of UMI outliers were removed with coverage at distant mapping locations on the DENV-1 genome.

### DENV genomic assemblies

Viral RNA was extracted from serum using Qiagen Viral RNA Mini QIAcube kit Qiagen Sciences, MD). Amplification was performed using 48 pairs of specific DENV-1 primers on the Fluidigm Access Array (Fluidigm Corporation, CA) with SuperScript III™ One-Step RT-PCR utilizing Platinum Taq High Fidelity polymerase (Thermo Fisher Scientific, MA). NGS libraries were prepared with QIAseq FX DNA Library Kit and sequenced on a Miseq with a v3 600 cycle sequencing kit. The sample reads were mapped to a DENV-1 Thailand strain (HM469967) using the NGS-mapper pipeline, that employs read quality filtering, trimming, mapping with BWA-MEM, and outputs assembly statistics and visualization for base call verification [54]. A minimum base quality threshold of 30.0 was used for base calling and variants were analyzed down to a 1% allele frequency threshold. Using NGS-mapper files, the consensus sequences were manually curated for accuracy.

The constructed consensus DENV-1 genomes from the serum samples were used as a reference in the cellranger pipeline. The resulting DENV-1 mapped BAM file, in conjunction with the cell barcodes obtained from cellranger count were used with 10x Geneomics subset-bam to exclude all reads not belonging to viable cells. Reads from the cell only BAM files were extracted using bedtools bamToFastq, and UMI-tools dedup was used to deduplicate UMIs. Optical duplicates were removed using BBMap clumpify and the reads were remapped with NGS-mapper using the serum genomes as reference and manually curated. A minimum base quality threshold of 25.0 was used for base calling and a 10% threshold was used for reporting all cell-associated genomic variants.

## Acknowledgments

The following reagent was obtained through the NIH AIDS Reagent Program, Division of AIDS, NIAID, NIH: CEM.NK^R^ Cells from Dr. Peter Cresswell.

## Data availability

The authors declare that all data supporting the findings of this study are available within this article or from the corresponding author upon reasonable request. Single-cell RNAseq gene expression data have been deposited in the Gene Expression Omnibus database.

## Disclaimer

The opinions or assertions contained herein are the private views of the authors and are not to be construed as reflecting the official views of the US Army, the US Department of Defense, or the National Institutes of Health. Material has been reviewed by the Walter Reed Army Institute of Research. There is no objection to its presentation and/or publication. The investigators have adhered to the policies for protection of human subjects as prescribed in AR 70–25.

